# Intracellular Expression of a Fluorogenic DNA Aptamer Using Retron Eco2

**DOI:** 10.1101/2024.05.21.595248

**Authors:** Mahesh A. Vibhute, Corbin Machatzke, Saskia Krümpel, Malte Dirks, Katrin Bigler, Daniel Summerer, Hannes Mutschler

## Abstract

DNA aptamers are short, single-stranded DNA molecules that bind specifically to a range of targets such as proteins, cells, and small molecules. Typically, they are utilized in the development of therapeutic agents, diagnostics, drug delivery systems, and biosensors. Although aptamers perform well in controlled extracellular environments, their intracellular use has been less explored due to challenges of expressing them in vivo. In this study, we employed the bacterial retron system Eco2, to express a DNA light-up aptamer in *Escherichia coli*. Our data confirms that structure-guided insertion of the aptamer domain into the non-coding region of the retron enables reverse transcription and biosynthesis of functional aptamer constructs in bacteria. The purified DNA aptamer synthesized under intracellular conditions shows comparable activity to a chemically synthesized control. Our findings demonstrate that retrons can be used to express short DNA aptamers within living cells, potentially broadening and optimizing their application in intracellular settings.

## Introduction

Aptamers are single stranded nucleic acids that fold into distinct structures, allowing them to bind specifically to target ligands.^1^ The ligand-binding sites of aptamers are tailored to the shape and charge of the ligand, enabling precise interactions. In particular, aptamers isolated through in vitro selection protocols serve as affinity reagents, positioning them as nucleic acid analogues of antibodies.^2,3^ As such, they offer distinct advantages over their protein-based counterparts such as small size, as well as rapid and cost-effective manufacturing.^3^ An area of particular interest is the development of fluorescent light-up aptamers (FLAPs),^4–6^ which specifically bind conditionally fluorescent dyes and activate their fluorescence.^7^ The advent of FLAPs has also significantly advanced the study of folding of aptamers, as efficient folding can be determined as a function of fluorescence output. Pioneering work by Jaffery and coworkers has led to proliferation of numerous FLAPs. Notable examples include the RNA-FLAPs Spinach^8^, Broccoli^9^, Corn^10^, Mango^11^, Squash^12^ and Pepper^7^. These fluorogenic RNA aptamers serve as RNA mimics of fluorescent proteins and have transformed cellular imaging techniques to locate and understand RNA dynamics in vivo.

In contrast to RNA aptamers, DNA aptamers offer enhanced stability, greater ease of synthesis and chemical modification, as well as reduced cost, thereby facilitating more straightforward production and broader applicability in diverse fields, including therapeutics and diagnostics.^3,13^ For example Lettuce, a DNA FLAP that mimics the fluorescent properties of GFP, holds great promise for pathogen detection or as intracellular sensors for tumour cell recognition.^14–16^ However, since most DNA aptamer studies focus on their development and characterization in vitro, their potential for applications inside living cells remains largely unexplored. A particular challenge in this context is the difficulty of biosynthesising DNA aptamers in cells. Indeed, the ability to transcribe single-stranded RNA from DNA-based vectors has decisively contributed to the development of diverse RNA FLAPs for in-cell applications, such as the Broccoli aptamer^9^. Therefore, methods for the intracellular synthesis and functional studies of single-stranded DNA (ssDNA) aptamers are highly desirable. Only a few approaches for intracellular ssDNA synthesis have been reported, in all cases using exogenous systems such as phagemid vectors^17^, phages^18^, and eukaryotic retroviral reverse transcriptases^19–23^.

A more integrated, efficient, and potentially safer alternative to exogenous ssDNA expression systems could be the use of endogenous bacterial platforms. Retrons are prokaryotic genetic elements that are part of the bacterial phage defence machinery.^24–27^ Each retron typically consists of three components, namely a ncRNA transcript, an RT, and an effector protein. The ncRNA transcript adopts a specific secondary structure, nested between two repeat regions at its 5’ and 3’ ends that are complementary and hybridize (Figure 1). The RT recognizes the folded ncRNA and initiates reverse transcription from the 2’OH of a conserved guanosine, using the base-paired region as primer.^28^ The reverse transcription proceeds until about halfway along the ncRNA. Simultaneously, the RNA template is degraded by RNase H1, except for a stretch of approximately 5-10 RNA bases, which remain hybridized to the cDNA.^24^ This results in a hybrid RNA-DNA structure (RT-DNA), with the non-coding RNA and DNA elements *msr* and *msd*, respectively. *msr* and *msd* are covalently attached by a 2’-5’ linkage at the conserved guanosine and the base paired region close to the 5’ end of *msd* (3’ end of *msr*). This ability of retrons to synthesize abundant ssDNA in vivo has generated considerable interest in using them as an alternative to exogenously delivered DNA oligonucleotides in genome engineering and genome editing.^28–35^

**Figure 1.**
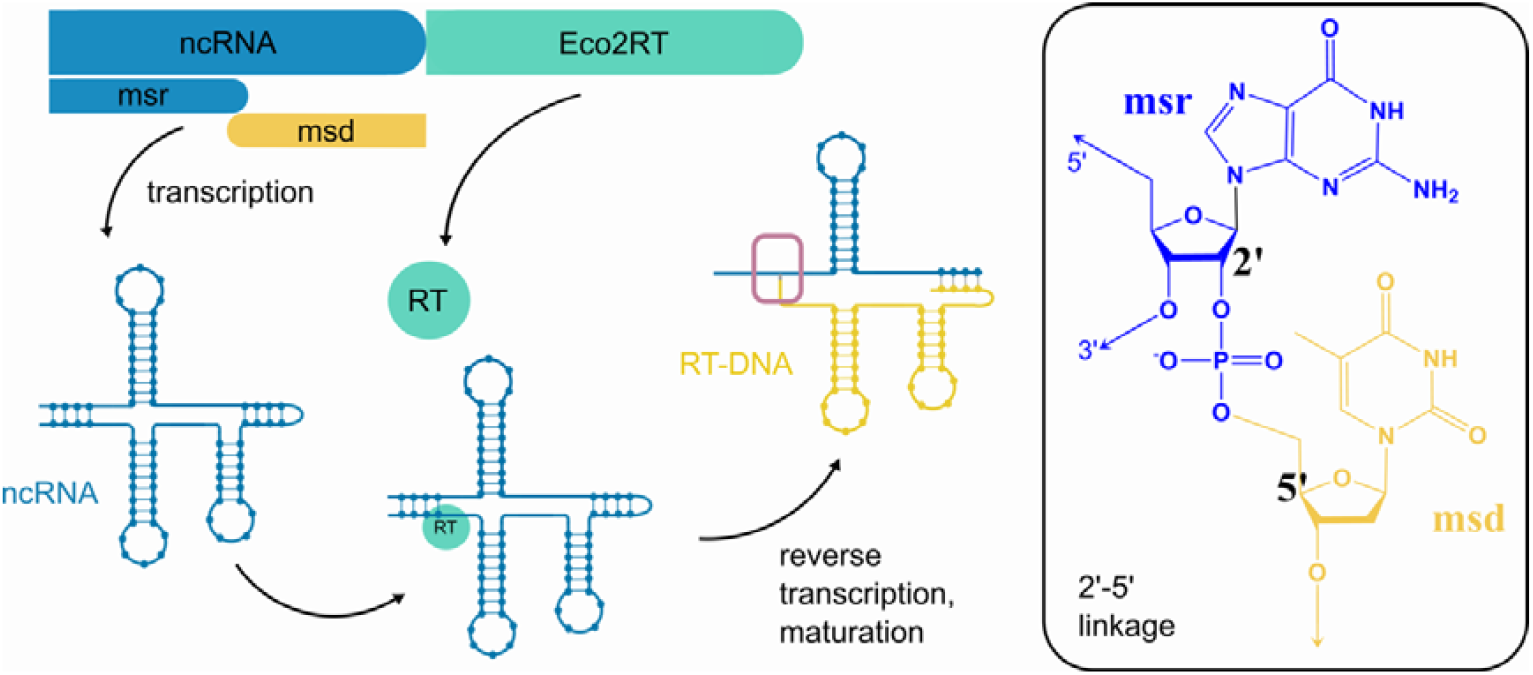
Schematic of the Eco2 gene locus, which encodes the non-coding RNA (ncRNA, blue) and the Eco2 reverse transcriptase gene (Eco2RT, green). The *msd* (yellow) region of the ncRNA is reverse transcribed into *msd* DNA, which is linked to the remaining *msr* via a 2’-5’ linkage conserved guanosine close to the 3’-end of the *msr*, as indicated in the inset.

While retrons have also been proposed as a potential method for expressing DNA aptamers,^20,22^ this approach has not yet been experimentally validated. Recent studies have indicated that retrons can indeed be leveraged to express functional ssDNA in vivo. For example, Lopez et al tested modifications to the ncRNA in case of retron Eco1 in order to boost RT-DNA abundance^32^. More recently, Liu et al also used retron Eco1 to express 10-23 DNAzyme in vivo^36^. However, it is poorly understood how the encoding of a functional ssDNA sequence in retrons will affect its activity. Such studies may lead to valuable design principles that would be especially important in the cellular context, as the RT-DNA is known to interact with not only the RT, but, depending on the retron, also RNaseH and the retron associated effector proteins.^24,25^

In this work we sought to demonstrate expression of a DNA FLAP Lettuce as a model using retron Eco2.^37^ We first characterize expression of plasmid-borne wild type Eco2, determining abundance and intracellular stability of Eco2 RT-DNA. Subsequently, we test suitable integration sites of different length variants of Lettuce within the retron *msd* using in vitro prototyping. After identifying effective insertion positions, we finally demonstrate fluorogenic activity of the expressed RT-DNA, embedded within the Lettuce aptamer. Our work demonstrates proof of concept of expressing DNA aptamers using retrons.

## Results

We first established an expression system for retron Eco2 by cloning the ncRNA and RT in a pET28a plasmid backbone, under the control of a T7 promoter. The plasmid was transformed in BL21AI cells, that enable tight control of T7-based expression. The intracellular synthesis of Eco2 RT-DNA expression was induced with 1 mM IPTG and 0,2% arabinose (Methods) and confirmed by denaturing PAGE of DNA extracts from induced cells. Using gel-based quantification of both RT-DNAs, we observed that Eco2 RT-DNA expression levels were approximately 3-4 times higher than those of genomically encoded Eco1 RT-DNA. To quantify expression levels, we further characterized Eco2 RT-DNA expression using qPCR. Upon induction with IPTG and arabinose, we observed that the copy-number ratio between amplicon 1 (which can arise from amplification of both single-stranded RT-DNA and plasmid DNA, as shown in Figure 2A) and amplicon 2 (which can only be amplified from the plasmid construct) increased by almost eightfold in induced compared to uninduced cells. Assuming an average copy number of 15-20 for the pET28b expression, which utilizes a pBR322 origin^38^, we estimate that 100-200 single-stranded RT-DNA molecules are present in induced cells. With an average cytoplasmic volume of 7 × 10^-16^ L of an *E. coli* cell^39^, this corresponds to an intracellular concentration of approximately 250 – 500 nM.

**Figure 2.**
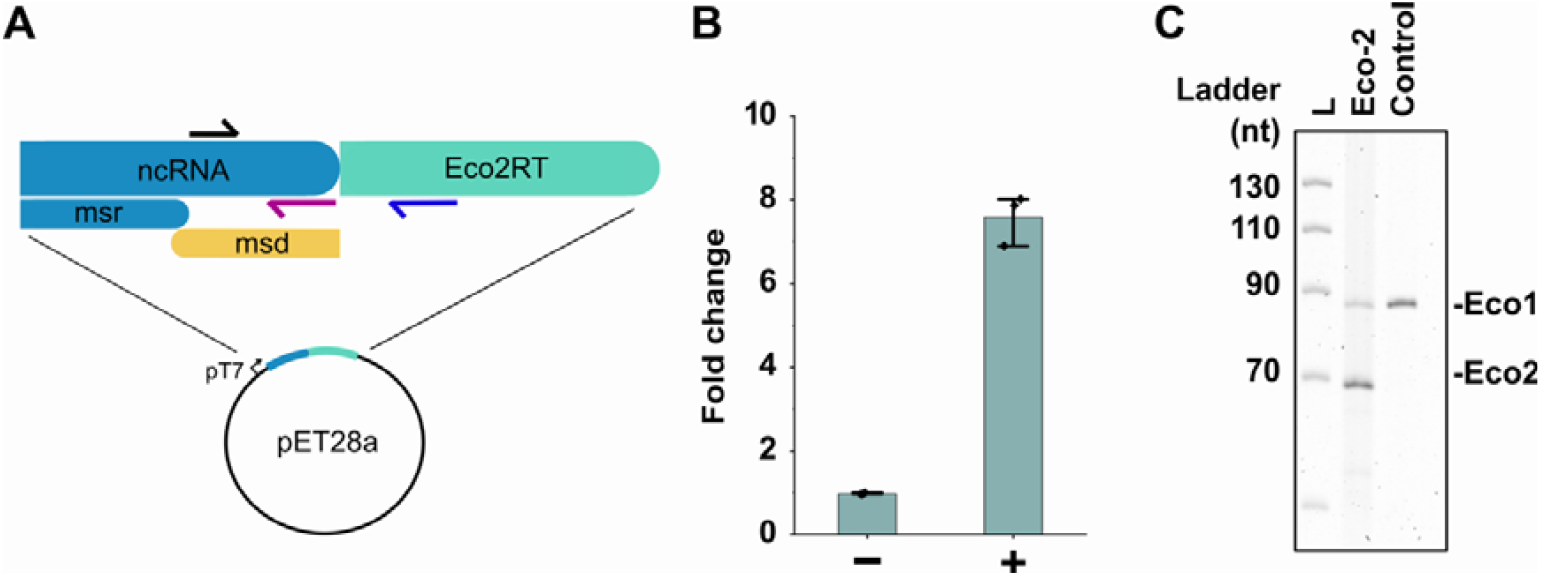
(A) Illustration of primer target sites for qRT-PCE experiments to determine RT-DNA abundance. The black arrows indicate the forward primer that pairs either with the purple reverse primer (amplify both RT-DNA and plasmid DNA) or the blue reverse primer that only amplified the plasmid DNA. (B) Fold enrichment of the RT-DNA/plasmid template over the plasmid alone upon induction, as measured by qPCR; Unpaired t-test, induced versus uninduced: p < 0.0001; n = 3 biological replicates. (C) A TBE-Urea polyacrylamide gel, stained with SYBR Gold showing RT-DNA corresponding to retron Eco1 (90 nt) and retron Eco2 (70 nt).

Because no data are available on the stability of single-stranded RT-DNA in cells, we next determined the half-life of Eco2 RT-DNA (Supplementary Information). To this end, induced cells were thoroughly washed to remove residual IPTG and arabinose, thereby preventing de novo expression of RT-DNA. The washed cells were then resuspended in fresh medium and allowed to continue growing. RT-DNA levels were quantified by qPCR at defined time points. The gradual decrease in RT-DNA abundance could thus be attributed to either active degradation (e.g. by intracellular nucleases) or dilution due to cell growth and division. Intriguingly, no evidence for active RT-DNA degradation was observed as the decrease in RT-DNA could be entirely explained by dilution (Figure 2–figure supplement 1) indicating exceptional intracellular stability.

Having established the inducible expression of plasmid-encoded Eco2 in *E. coli*, we next sought to explore its suitability as a scaffold to host a functional Lettuce aptamer. Expression of ssDNA using retrons is typically achieved by encoding a cargo gene in the *msd* region of the ncRNA. This approach has been successfully demonstrated for recombineering donors^32^, protein-binding DNA sequences^40^ and DNAzymes^36^. However, it was not clear as to what extent the folding of the cargo aptamer sequence would be affected due to the extensive secondary structure of the Eco2 *msr-msd* RNA-DNA hybrid.

We speculated that the position of the Lettuce aptamer domain in the *msd* region would significantly affect its ability to fold into the functional three-dimensional structure. To test this hypothesis, we used a structure-guided approach to insert different variants of the Lettuce aptamer as cargo into four different positions in the *msd* region of retron Eco2 (Figure 3). To identify four suitable insertion sites in the *msd*, we used the minimal free energy structure predicted for the Eco2 wild type *msd* by RNAfold^41^ with energy parameters for DNA as guide to identify single stranded and / or loop regions where an insertion of the Lettuce domain was expected to not interfere with the native fold of the remaining *msd* sequence. We omitted the *msr*-overlapping region in the input sequences as it is known to form a duplex with the RT-DNA as well as the *msr* RNA, as to the best of our knowledge, there is no algorithm capable of simulating mixed RNA-DNA sequences that are connected via a 2’-5’ bond. The positions v1 and v3 are in two different predicted loop regions. The insertions site v2 is located in a single stranded that connects two hairpin loops, whereas position v4 directly upstream of the predicted base-paired region between the *msd* and *msr* sequences.

**Figure 3.**
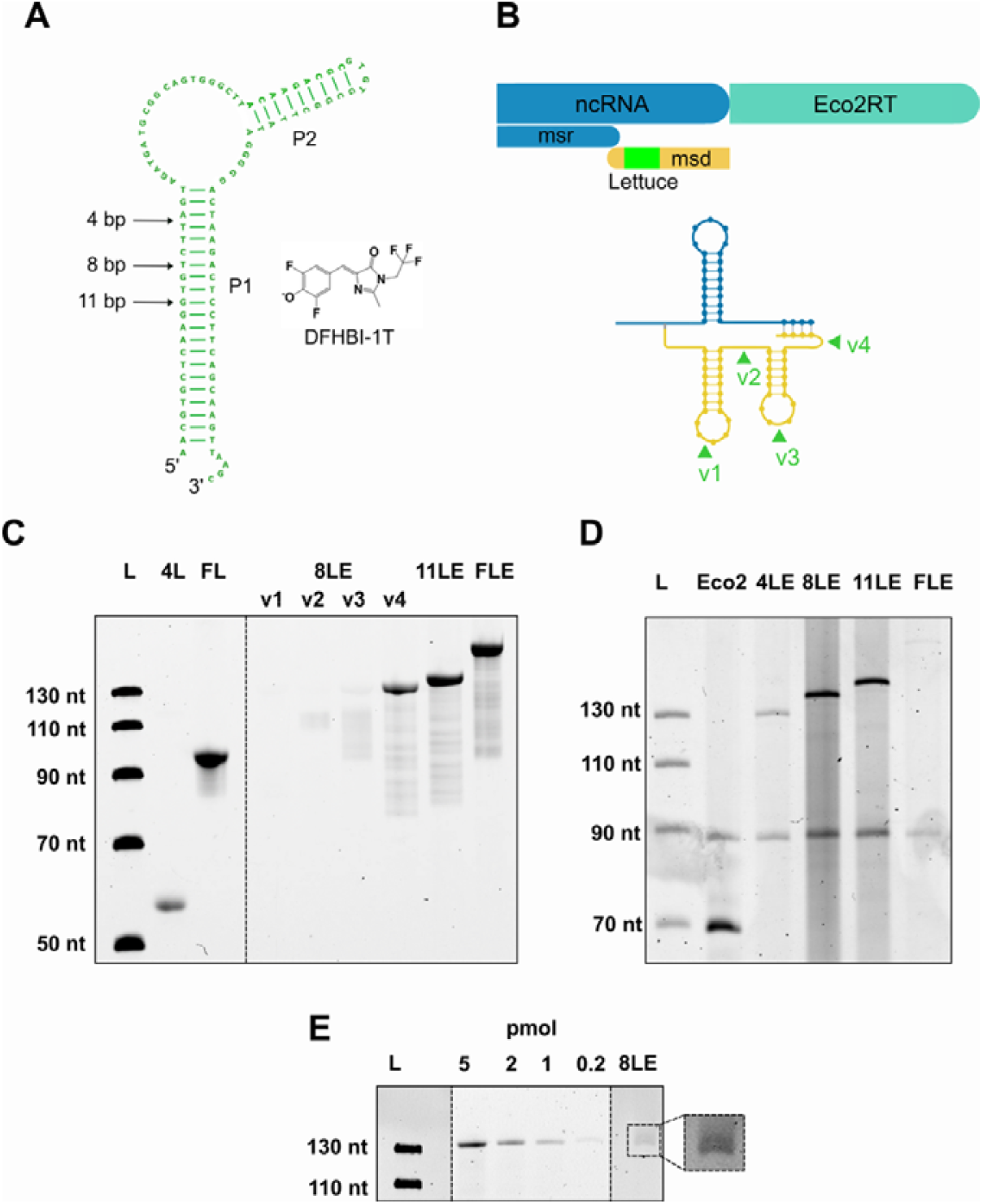
(A) Schematic of Lettuce aptamer. The arrows indicate the length to which the P1 stem can be shortened without significant loss of FLAP functionality. (B) Schematic of insertion of Lettuce aptamer sequence in 4 distinct positions in the *msd* region of retron eco2. The ssDNA structure was simulated using Vienna RNA fold with DNA parameters. Lettuce aptamer sequence was inserted at single stranded and/or loop regions to minimise interference with the native fold of the *msd* sequence. (C) DFHBI-1T stained TBE urea-PAGE showing fluorescence of different oligonucleotides mimicking the Lettuce-Eco2 (LE) fusion constructs with varying P1 lengths: 8LE v1-v4 (134 nt), 11LE (140 nt), and FLE (166 nt). Free Lettuce with either the 4 nt P1 stem (4L) or full-length P1 (FL) served as positive controls. (D) SYBR stained TBE urea-PAGE of extracted Eco2 wild type and Eco2-Lettuce RT-DNA fusions after expression in E. coli. In all variants, the lettuce aptamer was inserted into position v4 of the msd scaffold. (E) DFHBI-1T stained TBE urea-PAGE showing fluorescence of 8LEv4 RT-DNA purified from E. coli cells. Lanes with different loading amounts in pmol of the synthetic RT-DNA standards are shown as comparison.

The P1 stem of the original lettuce aptamer can be truncated down to four base pairs without significantly affecting its fluorescence output upon DFHBI-1T binding.^14^ We made use of this property in order to minimize the impact of Lettuce insertion on RT-DNA maturation while maintaining the ligand binding activity of Lettuce. Initially, we probed if truncated aptamer variants with P1 stem-lengths of 4, 8, and 11 base pairs (designated 4L, 8L, and 11L) along with the full-length aptamer (FL) can still fold after embedding them into the integration sites of the emulated Eco2 RT-DNA.

Using chemically synthesized RT-DNA-Lettuce fusion constructs as model constructs, we first tested the fluorogenic potential of 4L after its insertion into the above-mentioned positions v1 to v4 of the Eco2 RT-DNA (resulting in the constructs 4LEv1 – 4LEv4) using in gel staining with DFHBI-1T (Figure 3–figure supplement 1). We observed only very low levels of in-gel fluorescence compared to a standard full-length Lettuce (FL) or 4L alone, indicating that 4L folding was hampered upon embedding in the retron *msd* scaffold. In contrast, we observed a pronounced fluorescence signal for 8L when it was inserted position v4 (Figure 3C), demonstrating that this insertion site was compatible with Lettuce folding given a sufficient length of P1. Similarly, we observed strong in-gel fluorescence signals for 11LEv4 and FLEv4 comparable with native full-length Lettuce.

To cross-validate these results, we determined the dissociation constants between DFHBI-1T and the chemically synthesized Eco2-Lettuce v4 surrogate constructs using the reported Lettuce binding buffer (40 mM HEPES pH 7.5, 100 mM KCl, and 1 mM MgCl_2_)^14^. When increasing concentrations of each oligonucleotide were incubated in binding buffer containing a constant concentration of DFHBI-1T, we obtained well-defined fluorescence responses for all v4 constructs (Figure 3–figure supplement 2). When fitting the binding isotherms with a quadratic association curve (Supplementary Methods) we obtained dissociation constants (K_d_) for 8LE, 11LE, FLE that were comparable to the unembedded full-length aptamer (FL) ∼0.12 µM (0.01, 2.46, 95% CI). Binding was weaker for the synthesized ssDNA mimic 4LE-v4 with a less well-defined K_d_ of ∼6 µM (4.92, 7.18, 95% CI), which was well in agreement with our in-gel fluorescence experiments.

Having confirmed that the v4 surrogate of 8LE, 11LE and FLE constructs maintained the light-up aptamer properties of the embedded Lettuce sequences, we investigated whether the different length variants could be expressed in *E. coli* after integration into the plasmid-borne *msd* region of retron Eco2. PAGE analysis of RT-DNA from induced *E. coli* BL21-AI expressing the Eco2-Lettuce fusion constructs confirmed complete synthesis of the 8LE (134 nt) and 11LE (140 nt) species, and low-level production of FLE (166 nt), despite their *msd*-encoded RT-DNAs being more than twice as long as the wild-type (67 nt). Overall, the insertion of the Lettuce aptamer domain into the *msd* region of retron Eco2 resulted in a decrease in RT-DNA level compared to wild-type Eco2: While the length-corrected intensity of wild-type Eco2 was 3-fold higher than that of the endogenous Eco1 band, the 4LE, 8LE, 11LE and FLE constructs showed between ∼25% (FLE) and ∼65-90% (4LE, 8LE, and 11LE) of the length-corrected relative band intensities (Figure 3).

Next, we tested whether heterologously expressed 8LEv4 RT-DNA retains its fluorogenic properties, as it remained unknown whether retron-derived aptamer-DNAs produced under intracellular conditions are functionally equivalent to chemically synthesized ssDNA oligonucleotides. Because column-based purification could not be scaled efficiently due to the large amounts of plasmid DNA suppressing column binding of the short ssDNAs, we developed a modified TRIzol-based extraction protocol optimized for ssDNA isolation while minimizing co-purification of plasmid DNA. Although the new method yielded considerably cleaner RT-DNA preparations, overall recovery was lower, necessitating the use of larger culture volumes to obtain sufficient material for analysis. The new extraction method was evaluated in pilot experiments for the 8LEv1-4 variants, and PAGE analysis of the isolated RT-DNA from pilot-scale experiments confirmed full-length synthesis of all four position variants (Figure 3–figure supplement 3). This approach was then applied to larger volumes for preparative extractions. TBE-urea PAGE analysis with DFHBI-1T staining indeed revealed that the purified, intracellular ssDNA retains fluorogenic activity upon DFHBI-1T binding.

While no measurable fluorescence was detected in induced cell cultures expressing 8LEv4 - aside from DFHBI-1T-independent expression-related autofluorescence (Figure 3–figure supplement 4), the intracellular production of a fluorogen-binding RT-DNA demonstrates that retron-encoded ssDNA can, in principle, be used to generate functional aptamers in a cellular environment. The limited in vivo signal likely reflects both the moderate RT-DNA abundance (≈100-200 molecules per cell) and the substantially lower fluorogenic efficiency of Lettuce compared with the RNA light-up aptamer Broccoli (∼100-fold, Figure 3–figure supplement 5), which is optimized for in vivo imaging of RNA^9^. Nevertheless, the detection of fluorogenic activity in purified material establishes a proof-of-principle that retron-derived ssDNA can be used to display DNA aptamers inside living cells.

## Discussion

Bacterial retrons such as Eco2 provide a versatile platform for intracellular reverse transcription-based synthesis of single-stranded DNA. To our knowledge, our present study is the first to show that functional DNA aptamers that recognize small molecule ligands can also be expressed via retrons. Expanding the repertoire of retrons toward DNA aptamer expression promises to significantly expand the repertoire of functional non-coding DNAs and enable the creation of new sensors and regulators with a wide range of intracellular applications. At the same time, retron platforms provide a direct way to tailor DNA aptamers specifically to intracellular conditions. This is particularly important as our data show that both the low expression levels in combination with the moderate fluorogenic activity of Lettuce provide ample room for, e.g. evolutionary improvement of retron-encoded DNA aptamer performance under intracellular conditions. We observed a significant decrease in Eco2 expression levels after integration of the Lettuce domain, suggesting that reverse transcription is hindered by the changes in the *msd* sequence.^32^

In Eco2, the RT is directly fused to a topoisomerase-primase-like (Toprim) effector domain, whose activity causes abortive infection^42,43^. Recent structural studies elucidated the structure of the Eco2 nucleoprotein complex and revealed that each RT-Toprim fusion protein forms a 1:1 complex with the 2’-5’ linked DNA-RNA hybrid that further assembles into a trimer^42^. Interestingly, while the *msr* portion of each complex bridges adjacent RT–Toprim monomers and is integral to the structural integrity of the complex, the electron density of the *msd* beyond the stem downstream of the branched 2’-5’ linkage remained largely undefined, suggesting a strong degree of flexibility. The proximal *msd* stem is proposed to sterically block the nuclease active site of the Toprim effector domain; upon activation by the phage-encoded ssDNA nuclease DenB from T4-related phages, this inhibition is relieved, enabling Toprim to cleave tRNA and thereby inducing translational shutdown.^42,43^ The electron density of the distal *msd* region was largely unresolved in the reported cyro-EM structure, indicating that it is not an integral structural element of Eco2 and likely tolerates mutations or insertions without preventing trimerization. In our study, we indeed observed no major growth defects upon insertion of the aptamer domain at positions v1 to v4 confirming the high degree of tolerance of the *msd* region distal to the 2’-5’ branch point. This suggests that alternative functional sequences could be similarly integrated into the distal *msd* segment, provided that flanking *msr* and *msd* regions do not disrupt folding. A major advantage of retron-based intracellular synthesis of ssDNA in vivo is its simplicity. The desired sequence can be cloned into the *msd* region of plasmid-borne or genomically encoded retron Eco2. The copy number of the aptamer can also, in principle, be tuned by inducer concentration, promoter strength and plasmid copy number. An ever-increasing number of DNA aptamers have been generated against a wide range of targets, from small molecules, macromolecules to whole cells.^3,44^ The retron system used in this work could enable the exploration of cellular applications for this growing repertoire of DNA aptamers. A further advantage of expressing DNA aptamers with retrons is the exceptional intracellular stability of the *msd*-derived ssDNA in the retron complex (Supplementary information, Figure 2–figure supplement 1). Finally, expression of related retron systems has been demonstrated in yeast^45^ and mammalian cells^32,46^, thus potentially widening the range of in vivo applications of DNA aptamers.

In conclusion, retron-based aptamer expression platforms present a promising avenue for the development of novel DNA-based applications. Future research aimed at optimizing intracellular synthesis may elucidate the conditions under which larger single-stranded DNAs can be synthesized at higher concentrations without compromising reverse transcription and may also identify the key factors influencing the optimal in vivo folding of cargo sequences.

## Methods

### Constructs and strains

The retron Eco2 sequence (*msr*-*msd* and RT) was obtained from Millman et al^26^, and purchased from IDT as gblock, which was inserted into a pET28a vector under control of T7 promoter and lac operator. Plasmid assembly was performed using the HiFi DNA Assembly (New England Biolabs) according to the supplier’s instructions. The resulting plasmids were transformed into chemically competent *E. coli* Top 10 cells. Plasmid sequences were verified by colony PCR and Sanger sequencing (Microsynth AG). Verified plasmids were then transformed into chemically competent BL21AI cells for further experiments. These cells harbor endogenous retron Eco1 which serves as an internal control and a cassette encoding T7 polymerase driven by arabinose-inducible araBAD promoter. Lettuce variant sequences were cloned into retron Eco2 plasmid using Q5® Site-Directed Mutagenesis Kit (New England Biolabs). Primer sequences and annealing temperatures were generated using the NEBaseChanger™ tool.

### RT-DNA extraction

#### Miniprep method

5 ml cultures of BL21AI cells with corresponding Lettuce constructs (LB with selection antibiotic) were inoculated with cells from a single colony and grown at 37°C, 220 rpm for 90 min. They were then induced with 1 mM IPTG and 0.2% arabinose, followed by overnight expression. OD_600_ of the overnight culture was measured and cells corresponding to 2 ml of OD_600_ = 1 were used for miniprep. A NEB Monarch® Plasmid Miniprep Kit was used for RT-DNA extraction (New England Biolabs). Following miniprep, cells were eluted in 40 µl milliQ purified water. The miniprep product was then treated with a RNase Cocktail™ Enzyme Mix (Invitrogen) at 37°C for 30 min. The RNase treated samples were then purified using a ssDNA/RNA Clean & Concentrator kit (Zymo).

The purified RT-DNA was analysed on a 10% TBE-Urea polyacrylamide gel with a 1x TBE running buffer. Gels were stained with SYBR Gold (Invitrogen) and imaged on Sapphire Biomolecular Imager (Azure biosystems). Band intensities were normalized by length of RT-DNA. RT-DNA abundance of Eco2 and Lettuce variants was quantified by determining fold-change compared to the Eco1 band. Band intensity analysis was performed with ImageJ^47^.

#### TRIzol based extraction method

25 mL pre-cultures of BL21A cells transformed with the pET28a plasmid carrying the corresponding Lettuce variant were seeded from a single colony and grown in LB-medium containing 50 µg/mL kanamycin at 37 °C at 220 rpm for four hours until to an of OD_600_ of 0.5-0.8. These were used to inoculate 750 mL cultures, which were then grown at 37°C and 150 rpm for two hours. The cultures were then induced with IPTG (1 mM) and arabinose (0.2%) and allowed to grow overnight (∼16 h) until they reached an OD_600_ 2.4 - 2.9 and harvested by centrifugation at 4°C and 6000 g for 30 min. The cell pellets were then resuspended in TES buffer and lysozyme (Ready-Lyse Lysozyme Solution, Biozym) was added according to the manufacturer’s instructions (25 ml TES + 4.5 µL 30,000 U/µL Lysozyme). The samples were then incubated for 15 minutes at room temperature before they were split into 12 x 50 mL falcon tubes. 20 mL of TRIzol was added to each tube and the solutions were gently vortexed until they became clear. Next, 4 mL chloroform was added and the samples were vortexed until the solutions were homogeneous. The samples were then incubated at room temperature for 5 min until phase separation was observed. Subsequently, the samples were centrifuged for 30 min at 4°C and 12,000 g, after which a volume of 10 ml of the supernatant was added to 10 ml isopropanol. The samples were vortexed, then incubated for two hours at −20 °C. They were then centrifuged at 30,000 g and 4°C for 1 hour. The supernatant was then removed, and the pellet was washed twice with 10 ml of 70% ethanol and centrifuged at 4 °C for 5 minutes at 10,000 g. The pellets were air-dried for 15 minutes and then resuspended in 1000 μL of ddH_2_O.

### In-gel staining of the RT-DNA

After isolation using the above-described TRIzol-based method, 650 µL of the total RNA / retron (1000 µL) material was incubated with 250 mM NaOH and 6.25 mM EDTA at 80 °C for 1 hour. After incubation, the pH was neutralised by the addition of 250 mM HCl, after which the ssDNA was purified using the ssDNA/RNA Clean & Concentrator Kit (Zymo Research) following the manufacturer’s instructions. A total of three columns were used to isolate these samples, with 9 µL of ddH_2_O for elution. The eluate from column 1 was used to elute DNA from columns 2 and 3 to enhance the total RT-DNA concentration. The entire sample (approximately 7.5 µL), was mixed with 15 µL of loading dye (10 mM EDTA in formamide) and heated to 95 °C for three minutes, followed by immediate incubation on ice for five minutes. The sample was then loaded into a single well of a 10% TBE urea-PAGE gel and run for 80 minutes at a constant power of 20 W. The gel was then washed twice with ddH_2_O to remove the urea and EDTA. This was followed by a 30-minute stain with 50 mL DFHBI-1T staining solution, containing 40 mM HEPES pH 7.5, 100 mM KCl, 1 mM MgCl_2_ and 10 µM DFHBI-1T. After a final five-minute wash in ddH_2_O to remove excess dye, the gel was imaged at 488/518 nm (DFHBI-1T) and 520/565 nm (Cy3-tagged ladder) on a Sapphire Bioimager.

### qPCR

qPCR experiments were performed using Techne Prime Pro 48 Real-Time qPCR system, according to comparative C_T_ method,^48,49^ as described by Lopez et al.^32^ For qPCR analysis, cell cultures grown for ∼16h were used. 25 µl of bacterial culture at an OD_600_=1 was mixed with 25 µl water and incubated at 95°C, for 5 min to lyse cells. 0.3 µl of this boiled culture was used as templates in 30 µl qPCR reactions using Luna® Universal qPCR Master Mix (New England Biolabs). qPCR quantification of RT-DNA abundance was performed using a set of three primers. Two primers were complementary to the *msd* region of Retron Eco2 and the third was an outnest primer complementary to non-retron elements of the expression plasmid. Two templates were thus amplified: One corresponding to an RT-DNA sequence that could use both the RT-DNA and the plasmid as templates, and a second that could use only the plasmid-encoded Eco2 as a template. The difference between cycle threshold (ΔC_T_) of the inside (RT-DNA) and outside (plasmid) primer sets was then calculated. This ΔC_T_ was then subtracted from the ΔC_T_ of uninduced samples. Fold enrichment of RT-DNA/plasmid over plasmid alone was then calculated as 2^-ΔΔC^^T^, for each biological replicate. Presence of RT-DNA results in a fold change > 1.

## Competing Interest Statement

The authors declare no competing interest.

## Author Contributions

M.V., C.M., D.S., H.M. designed research; M.V., C.M., S.K., M.D. performed research; M.V., C.M., K.B., H.M., analysed data; K.B. contributed new reagents/analytic tools; M.V., C.M. and H.M wrote the paper.

## Acknowledgements

The authors would like to thank Indrayani Phadtare and members of Mutschler lab for fruitful discussions. HM acknowledges support by the European Research Council (ERC) under the Horizon 2020 research and innovation program (grant agreement ID: 802000, RiboLife).

## Notes

### Competing Interest Statement

The authors have declared no competing interest.

### Summary of Updates

All Figures revised. Text updated. Author list updated, Supplemental files updated. Major changes: -Cellular fluorescence section revised to account for stress-induced autofluorescence and to remove the previous in vivo Lettuce fluorescence claim. -Added in vivo stability measurements for Eco2 RT-DNA -Added new in vitro DFHB-1T affinity measurements for Eco2-Lettuce constructs -Added benchmarking of Lettuce vs Broccoli fluorogenicity -Figure 3 and associated supplements updated with new ex vivo data added demonstrating functional fluorescence of retron-produced Lettuce RT-DNA after purification from E. coli.

