## Supplementary Information for "Intracellular Expression of a Fluorogenic DNA Aptamer Using Retron Eco2"

Mahesh A. Vibhute#, Corbin Machatzke#, Saskia Krümpel, Malte Dirks, Katrin Bigler, Daniel Summerer and Hannes Mutschler*
Department of Chemistry and Chemical Biology, TU Dortmund University, Otto-Hahn-Str. 4a, 44227 Dortmund

#contributed equally

### Supplementary methods

#### Dissociation constant measurements

Dissociation constant measurements were performed on a BMG Labtech Clariostar plus plate reader. Oligonucleotides corresponding to different variants of Lettuce embedded in the retron scaffold were synthesized by IDT. Sample fluorescence was measured in a a Greiner 384-well black-bottom plate with excitation at 470±8 nm and emission at 515±20 nm. The samples were prepared by mixing 40 mM HEPES pH 7.4, 100 mM KCl, 1 mM $MgCl_2$ with different concentrations of DNA. Samples were then heated to 90 °C for 2 minutes, followed by incubating at room temperature for 5 minutes. After that 1 µM of DFHBI-1T was added to the mix. The samples were then briefly vortexed and spun down, and pipetted in triplicate into a 384-well plate. Samples were then incubated for 60 minutes at RT in darkness, followed by fluorescence measurements on the microplate reader.

The datasets were fitted using equation (1)

$$F = C_1 \cdot \frac{(A_t + B_t + k_D) - \sqrt{(A_t + B_t + k_D)^2 - 4 \cdot (A_t \cdot B_t)}}{2 \cdot B_t} + C_0 \qquad (1)$$

where F is the measured fluorescence, $A_t$ is the concentration of DNA, $B_t$ is the concentration of DFHBI-1T and $C_0$ and $C_1$ are the lower and upper plateau. To compensate for imprecision in determining the precise DFHBI-1T stock concentration, $B_t$ was also fit using strong constraints. Curve fitting was performed using the least_squares function from the *scipy.optimize* package (https://docs.scipy.org/doc/scipy/reference/optimize.html) with shared global parameters $C_0$, $C_1$ and $B_t$ for all data series.

#### Determination of half-life of Eco2 RT-DNA

Retron RT-DNA forms a phage surveillance complex with the associated RT and effector protein.[1–4] Owing to its unique 'closed' structure[5] (with the ends of *msr* and *msd* joined by a 2'-5' linkage and a base-paired region) and its non-coding function, the intracellular stability of RT-DNA is of particular interest. To assess the stability of Eco2 RT-DNA, we determined its half-life by qPCR.

We first induced retron Eco2 RT-DNA expression in BL21AI cells with 1 mM IPTG and 0.2% arabinose overnight, as described in the Methods section in the main text. On the following day, cells were pelleted and washed twice with LB medium to remove residual IPTG and arabinose. The washed cells were then resuspended and used to inoculate a fresh culture in LB medium without inducers at an initial $OD_{600}$ of 0.2. The culture was grown at 37 °C, and aliquots were taken at defined timepoints. For each time point, a cell suspension with $OD_{600}$ = 0.1 was prepared and incubated at 95°C for 5 min. One microliter of this boiled, diluted culture was used as input for a 20 µ l qPCR reaction. qPCR experiments were performed as described in the Methods section.

Assuming RT-DNA degradation would occur by active degradation mechanisms (nuclease mediated degradation) and dilution (cell growth and division), we determined the rate of degradation by the following equation

$$C(t) = 1 + A \cdot e^{-kt} \frac{OD_{600}(t_0)}{OD_{600}(t)} \qquad (1)$$

where $k$ is the degradation rate constant and the ratio $\frac{OD_{600}(t_0)}{OD_{600}(t)}$ is the dilution factor which takes into account dilution by cell division. $OD_{600}(t)$ was determined by fitting the $OD_{600}$ values to the following equation describing logistic growth (Figure S6A):

$$OD_{600}(t) = \frac{OD_{max}}{1 + e^{-r \cdot (t - t_0)}} \qquad (2)$$

After substituting $OD_{600}(t)$ with the function in equation (2), we fitted the experimental data for the fold-change of the RT-DNA to equation (1) (Figure S6B). Interestingly, the best fit (red) was obtained with a rate constant $k$ converging towards zero, suggesting that the half-life of the Eco 2 RT-DNA is beyond the detection limit of our assay. To illustrate typical half-lives of RNA, which are on the order of minutes in growing *E. coli* cells[6], we refitted the data using constant half-lives of 15 and 30 minutes. In both cases, the best-fit curves deviated substantially from the experimental data further supporting that the half-life of the RT-DNA is probably orders of magnitude longer than the doubling time of *E. coli* under these optimal growth conditions. While we cannot exclude that the RT-DNA is still produced due to promoter leakiness, we expect this effect to be minor, as the expression of RT-DNA requires the presence of both IPTG and arabinose, which were thoroughly removed before inoculating the growth medium with the starter culture. Overall, our data support the notion of exceptional RT-DNA stability.

#### In vivo fluorescence measurements

5 ml cultures (LB with selection antibiotic) were seeded with a colony and grown at 37°C, 220 rpm for 90 min in a 10 ml culture tube. They were then induced with 1 mM IPTG and 0.2% arabinose, followed by growth for ~16 h. A culture volume corresponding to 5 ml at $OD_{600}$ = 1 was pelleted and washed twice with 1 ml 1x PBS. The pellet was then resuspended in 150 µL 1x PBS and 40 µM DFHBI-1T was added to the cell suspension, followed by incubation at room temperature for 20 min. The stained cells were then used for plate reader measurements (3 x 50 µL in a 96-well plate). Plate reader measurements were performed with a BMG Labtech Clariostar plus plate reader at excitation and emission wavelengths of 470 nm and 520 nm, respectively. To normalise for variance in pipetting, we measured the $OD_{600}$ of the cells in 1x PBS after taking the microplate reader measurements. We used a 25-fold dilution on a Nanodrop to take these measurements, and then used the results to normalise the plate reader measurements.

#### Supplementary Figures

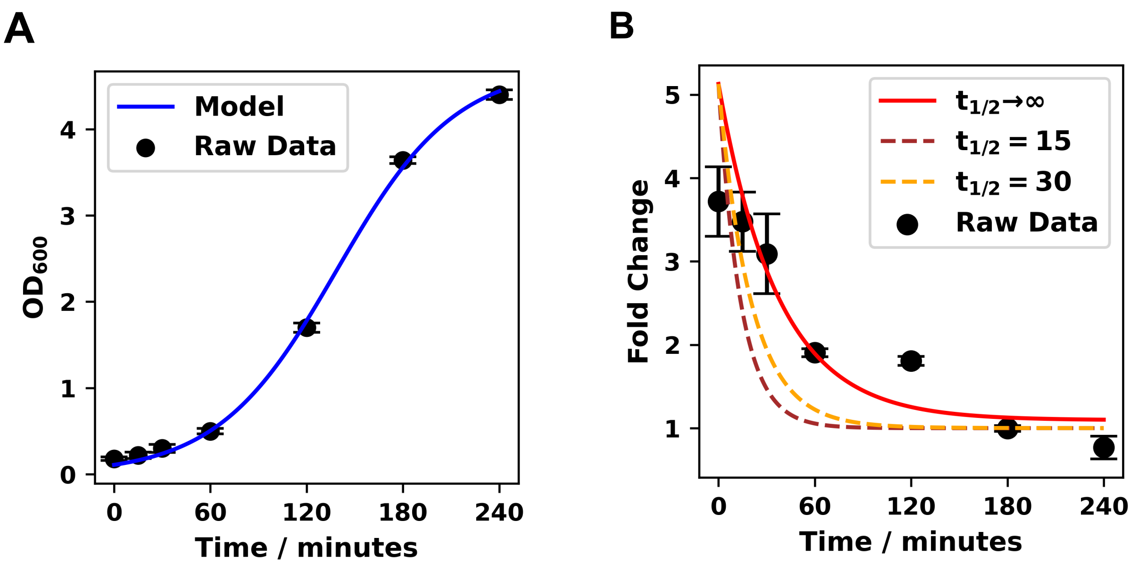

**Figure 2–figure supplement 1**. Determination of half-life of RT-DNA. (A) $OD_{600}$ measurements fitted with the equation for logistic growth. (B) Fold change of RT-DNA/plasmid amplicon over plasmid alone, at different time points. The data corresponds to n = 5 biological replicates with error bars indicating s.e.m.

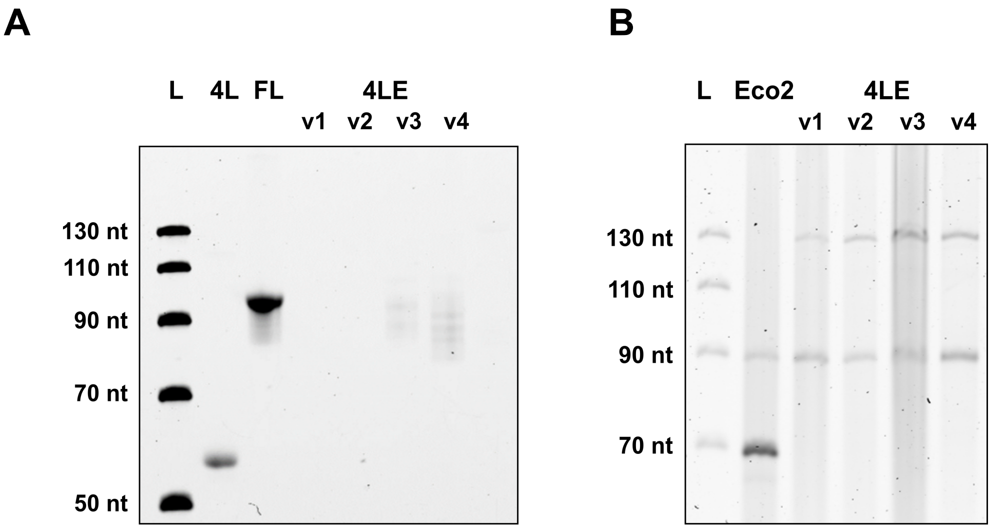

**Figure 3–figure supplement 1.** (A) In-gel DFBI-1T staining of 4LE oligonucleotides mimicking the 4Lettuce length variants in Eco2, with free 4Lettuce and full Lettuce aptamer as positive controls. (B) SYBR-stained gel showing expression of 4LEv1-4.

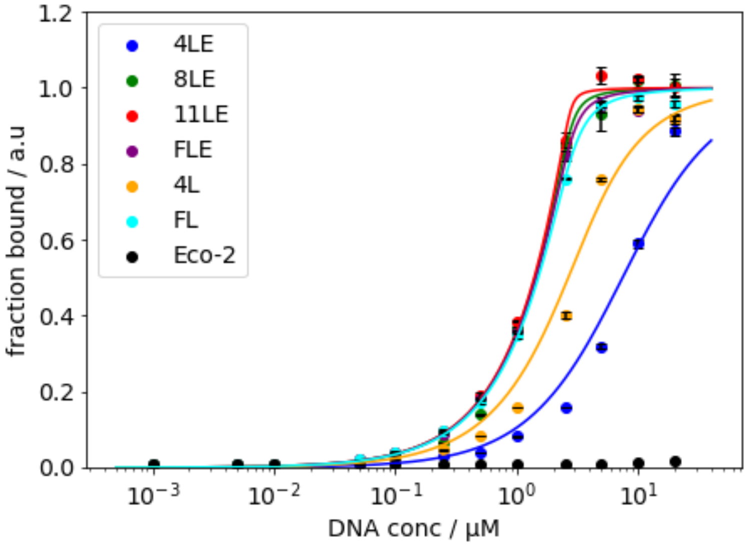

**Figure 3–figure supplement 2**. Fluorescence titration experiments using chemically synthesized Lettuce-Eco2 v4 variants. Solid lines represent plots of equation (1) using the best fit parameters. Fluorescence was measured at a fixed concentration of 1 μM DFHBI-1T and increasing DNA concentrations. Experimental data points are averages from n = 3 biological replicates with error bars indicating standard deviations. Calculated $K_D$s are: 4LE-v4 6 μM (4.93, 7.18, 95% CI); 8LE-v4 0.04 μM (0.01, 0.12, 95% CI); 11LE-v4 0.01 μM (0.01, 0.07, 95% CI); FLE-v4 0.07 μM (0.01, 0.16, 95% CI); 4L 1.28 μM (0.9, 1.6, 95% CI); FL 0.12 μM (0.01, 0.25, 95% CI).

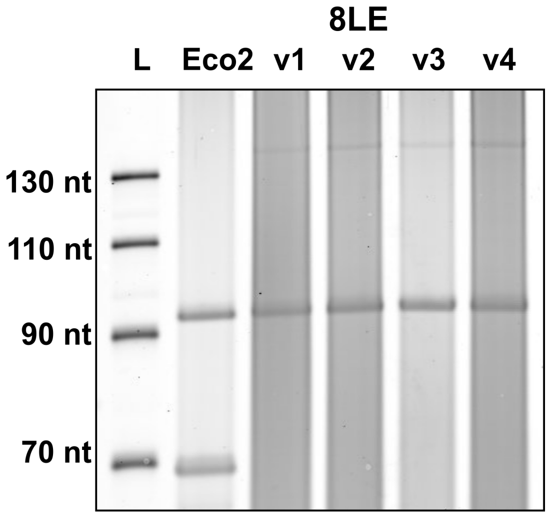

**Figure 3–figure supplement 3**. SYBR-stained TBE urea-PAGE showing TRIzol extracted RT-DNA corresponding to Eco2 (90 nt) and 8LEv1-4 (134 nt). The total extracted RNA was quantified using Qubit and an amount corresponding to 2000 ng RNA was loaded on the gel.

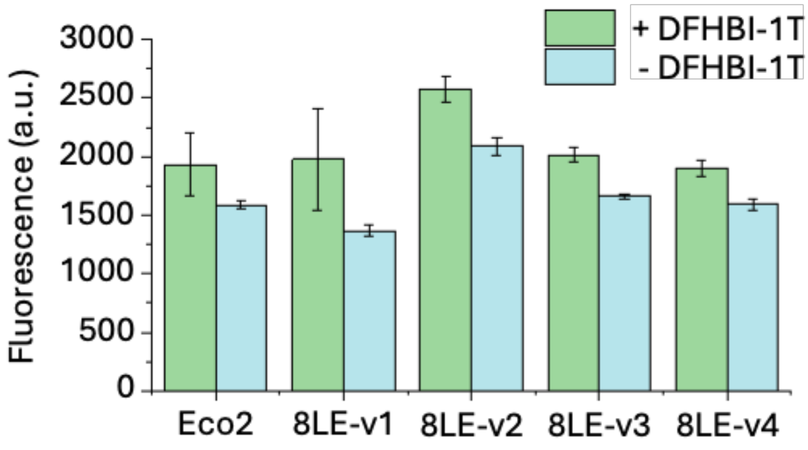

**Figure 3–figure supplement 4.** Bulk in vivo fluorescence measurements of 8LEv1-4 with DFHBI-1T. Experimental data points are averages from n = 3 biological replicates +/- standard deviations.

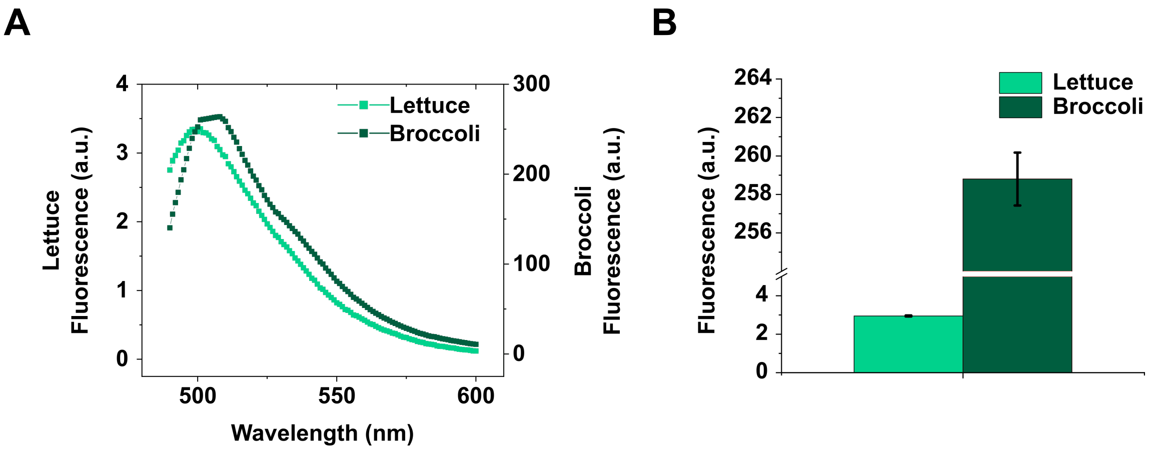

**Figure 3–figure supplement 5**. Comparison of fluorescence of Lettuce (DNA) and Broccoli (RNA) aptamers. (A) Emission spectra of equivalent concentrations (10 µM) of DNA Lettuce aptamer and RNA Broccoli aptamer, with equimolar DFHBI-1T, upon excitation at 462 nm. (B) Fluorescence output of Lettuce and Broccoli aptamer upon excitation at 462 nm, recorded at 510 nm.

### Sequences

| Table S1 | |
|---|---|
| Retron Eco2 | CATAAACACGCATGTAGGCAGATTTGTTGGTTGTGAATCGCAACCAGTGGCCTTAATGGCAGGAGGAATCGC<br>CTCCCTAAAATCCTTGATTCAGAGCTATACGGCAGGTGTGCTGTGCGAAGGAGTGCCTGCATGCGTTTCTCCT<br>TGGCCTTTTTTCCTCTGGGATGAAGAAGAAATGACAAAAACATCTAAACTTGACGCACTTAGGGCTGCTACTT<br>CACGTGAAGACTTGGCTAAAATTTTAGATgTTAAGTTGGTATTTTTAACTAACGTTCTATATAGAATCGGCTCGG<br>ATAATCAATACACTCAATTTACAATACCGAAGAAAGGAAAAGGGGTAAGGACTATTTCTGCACCTACAGACCG<br>GTTGAAGGACATCCAACGAAGAATATGTGACTTACTTTCTGATTGTAGAGATGAGATCTTTGCTATAAGGAAA<br>ATTAGTAACAACTATTCCTTTGGTTTTGAGAGGGGAAAATCAATCATCCTAAATGCTTATAAGCATAGAGGCAA<br>ACAAATAATATTAAATATAGATCTTAAGGATTTTTTTGAAAGCTTTAATTTTGGACGAGTTAGAGGATATTTTCT<br>TTCCAATCAGGATTTTTTATTAAATCCTGTGGTGGCAACGACACTTGCAAAAGCTGCATGCTATAATGGAACCC<br>TCCCCCAAGGAAGTCCATGTTCTCCTATTATCTCAAATCTAATTTGCAATATTATGGATATGAGATTAGCTAAGCT<br>GGCTAAAAAATATGGATGTACTTATAGCAGATATGCTGATGATATAACAATTTCTACAAATAAAAATACATTTCCG<br>TTAGAAATGGCTACTGTGCAACCTGAAGGGGTTGTTTTGGGAAAAGTTTTGGTAAAAGAAATAGAAAACTCT<br>GGATTCGAAATAAATGATTCAAAGACTAGGCTTACGTATAAGACATCAAGGCAAGAAGTAACGGGACTTACA<br>GTTAACAGAATCGTTAATATTGATAGGTGTTATTATAAAAAAACTCGGGCGTTGGCACATGCTTTGTATCGTACA<br>GGTGAATATAAAGTGCCAGATGAAAATGGTGTTTTAGTTTCAGGAGGTCTGGATAAACTTGAGGGGATGTTT<br>GGTTTTATTGATCAAGTTGATAAGTTTAACAATATAAAGAAAAAACTGAACAAGCAACCTGATAGATATGTATT<br>GACTAATGCGACTTTGCATGGTTTTAAATTAAAGTTGAATGCGCGAGAAAAAGCATATAGTAAATTTATTTACT<br>ATAAATTTTTTCATGGCAACACCTGTCCTACGATAATTACAGAAGGGAAGACTGATCGGATATATTTGAAGGCT<br>GCTTTGCATTCTTTGGAGACATCATATCCTGAGTTGTTTAGAGAAAAAACAGATAGTAAAAAGAAAGAAATAA<br>ATCTTAATATATTTAAATCTAATGAAAAGACCAAATATTTTTTAGATCTTTCTGGGGGAACTGCAGATCTGAAAA<br>AATTTGTAGAGCGTTATAAAAATAATTATGCTTCTTATTATGGTTCTGTTCCAAAACAGCCAGTGATTATGGTTCT<br>TGATAATGATACAGGTCCAAGCGATTTACTTAATTTTCTGCGCAATAAAGTTAAAAGCTGCCCAGACGATGTAA<br>CTGAAATGAGAAAGATGAAATATATTCATGTTTTCTATAATTTATATATAGTTCTCACACCATTGAGTCCTTCCGG<br>CGAACAAACTTCAATGGAGGATCTTTTCCCTAAAGATATTTTAGATATCAAGATTGATGGTAAGAAATTCAACA<br>AAAATAATGATGGAGACTCAAAAACGGAATATGGGAAGCATATTTTTTCCATGAGGGTTGTTAGAGATAAAAA<br>GCGGAAAATAGATTTTAAGGCATTTTGTTGTATTTTTGATGCTATAAAAGATATAAAGGAACATTATAAATTAAT<br>GTTAAATAGCTAATGAACAGCCCTAACGTTATGAACGCTAAGGCTGATTTTTCGTTAAAATTTATATGGTTTGAA<br>TTGTAATATATTATCTTCAAGCCATTTATTTAATTCCTGC |

Oligonucleotides / Ultramers used for in vitro measurements

| Table S2 | |
|---|---|
| 4LE-v1 | TCCTTCGCACAGCACATAGTAGGGATGATGCGGCAGTGGGCTTCGCAGAACAGTGTTTATTCTGCGAGGGGA<br>CTACCTGCCGTATAGCTCTGAATCAAGGATTTTAGGGAGGCGATTCCTCCTGCC |
| 4LE-v2 | TCCTTCGCACAGCACACCTGCCGTATTAGTAGGGATGATGCGGCAGTGGGCTTCGCAGAACAGTGTTTATTCT<br>GCGAGGGGACTAAGCTCTGAATCAAGGATTTTAGGGAGGCGATTCCTCCTGCC |
| 4LE-v3 | TCCTTCGCACAGCACACCTGCCGTATAGCTCTGAATCAATAGTAGGGATGATGCGGCAGTGGGCTTCGCAGAA<br>CAGTGTTTATTCTGCGAGGGGACTAGGATTTTAGGGAGGCGATTCCTCCTGCC |
| 4LE-v4 | TCCTTCGCACAGCACACCTGCCGTATAGCTCTGAATCAAGGATTTTAGGGAGGCGATTAGTAGGGATGATGCG<br>GCAGTGGGCTTCGCAGAACAGTGTTTATTCTGCGAGGGGACTATCCTCCTGCC |
| 8LE-v1 | TCCTTCGCACAGCACAGTCTTAGTAGGGATGATGCGGCAGTGGGCTTCGCAGAACAGTGTTTATTCTGCGAG<br>GGGACTAAGACCCTGCCGTATAGCTCTGAATCAAGGATTTTAGGGAGGCGATTCCTCCTGCC |
| 8LE-v2 | TCCTTCGCACAGCACACCTGCCGTATGTCTTAGTAGGGATGATGCGGCAGTGGGCTTCGCAGAACAGTGTTTA<br>TTCTGCGAGGGGACTAAGACAGCTCTGAATCAAGGATTTTAGGGAGGCGATTCCTCCTGCC |
| 8LE-v3 | TCCTTCGCACAGCACACCTGCCGTATAGCTCTGAATCAAGTCTTAGTAGGGATGATGCGGCAGTGGGCTTCGC<br>AGAACAGTGTTTATTCTGCGAGGGGACTAAGACGGATTTTAGGGAGGCGATTCCTCCTGCC |

| | |
|---|---|
| 8LE-v4 | TCCTTCGCACAGCACACCTGCCGTATAGCTCTGAATCAAGGATTTTAGGGAGGCGATGTCTTAGTAGGGATGATGCGGCAGTGGGCTTCGCAGAACAGTGTTTATTCTGCGAGGGGACTAAGACTCCTCCTGCC |
| 11LE | TCCTTCGCACAGCACACCTGCCGTATAGCTCTGAATCAAGGATTTTAGGGAGGCGATGGTGTCTTAGTAGGGATGATGCGGCAGTGGGCTTCGCAGAACAGTGTTTATTCTGCGAGGGGACTAAGACTCCTCCTCCTGCC |
| FLE | TCCTTCGCACAGCACACCTGCCGTATAGCTCTGAATCAAGGATTTTAGGGAGGCGATAACGTGCTCAAGGTGTCTTAGTAGGGATGATGCGGCAGTGGGCTTCGCAGAACAGTGTTTATTCTGCGAGGGGACTAAGACTCCTTCAGCAAGTTAAGCTCCTCCTGCC |

qPCR primers

| Table S3 | |
|---|---|
| Eco2 inside *msd* FWD | AGGAGGAATCGCCTCCCTAA |
| Eco2 inside *msd* REV | TCCTTCGCACAGCACACCT |
| outside *msd* REV | AAGGCCAAGGAGAAACGCAT |

Sanger sequencing primers

| Table S4 | |
|---|---|
| TdT Fill in P7 Adaptor: | GTGACTGGAGTTCAGACGTGTGCTCTTCCGATCTTTTTTTTTTTTTTTTTTTTTTTTTTTTTTTVN |
| 2nd Adaptor primer | ACACTCTTTCCCTACACGACGCTCTTCCGATCTTCCTTCGCACAGCACACCTGCCGTATAG |
| Fill in reverse primer | GTGACTGGAGTTCAGACGTGTGCTCTTCCGATCT |
